# Rapid Adeno-Associated Virus Genome Quantification with Amplification-Free CRISPR-Cas12a

**DOI:** 10.1101/2023.11.14.567134

**Authors:** Zach Hetzler, Stella M. Marinakos, Noah Lott, Noor Mohammad, Agnieszka Lass-Napiorkowska, Jenna Kolbe, Lauren Turrentine, Delaney Fields, Laurie Overton, Helena Marie, Angus Hucknall, Oliver Rammo, Henry George, Qingshan Wei

## Abstract

Efficient manufacturing of recombinant Adeno-Associated Viral (rAAV) vectors to meet rising clinical demand remains a major hurdle. One of the most significant challenges is the generation of large amounts of empty capsids without the therapeutic genome. There is no standardized analytical method to accurately quantify the viral genes, and subsequently the empty-to-full ratio, making the manufacturing challenges even more complex. We propose the use of CRISPR diagnostics (CRISPR-Dx) as a robust and rapid approach to determine AAV genome titers. We designed and developed the CRISPR-AAV Evaluation (CRAAVE) assay to maximize sensitivity, minimize time-to-result, and provide a potentially universal design for quantifying multiple transgene constructs encapsidated within different AAV serotypes. We also demonstrate an on-chip CRAAVE assay with lyophilized reagents to minimize end user assay input. The CRAAVE assay was able to detect AAV titers as low as 7e7 vg/mL with high precision (<3% error) in quantifying unknown AAV titers when compared with conventional quantitative PCR (qPCR) method. The assay only requires 30 minutes of assay time, shortening the analytical workflow drastically. Our results suggest CRISPR-Dx could be a promising tool for efficient rAAV genome titer quantification and has the potential to revolutionize biomanufacturing process analytical technology (PAT).

**Graphical Abstract:** 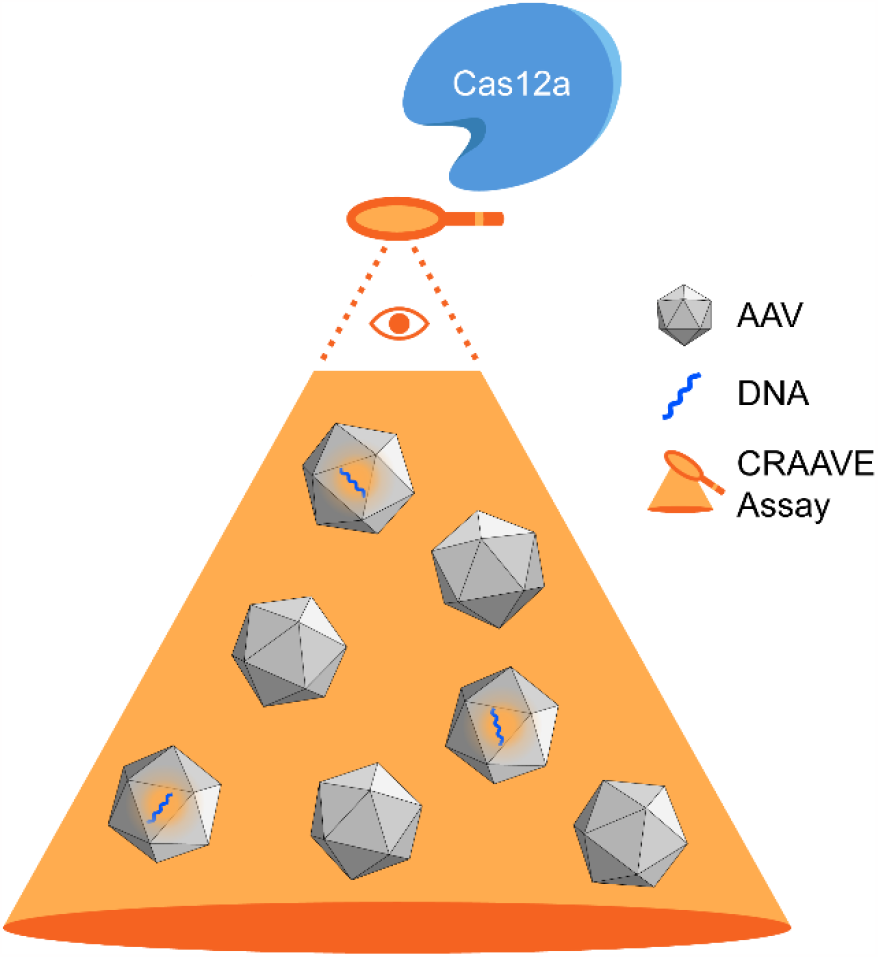

## Introduction

Adeno-Associated Virus (AAV) is a small, nonenveloped, replication incompetent virus containing a ssDNA genome of just under 5 kilobases. AAV was originally discovered as a contaminant in adenovirus cultures, leading to its similar name [1]. Recombinant adeno-associated virus (rAAV) has become one of the most promising gene therapy vectors due to its nonpathogenic property, well-established tissue tropism, and long-term transgene expression [2], [3]. There are over 100 active gene therapy clinical trials using AAV as the delivery vector [4], with many more AAV gene therapies in the preclinical pipeline. A primary challenge to meet this quickly growing clinical demand for AAV vectors is efficiently manufacturing large amounts of AAV with high potency. AAV industrial manufacturing is faced with several difficult problems such as generating high titers, formation of AAV aggregates, and developing a scalable transfection process. Perhaps the most problematic of all is the generation of a high level of empty capsids [5], a viral particle nearly identical to the desired product, distinguished only by the lack of an encapsidated therapeutic transgene. Definitions vary as to whether capsids containing partial, fragmented transgenes are considered empty capsids. Further muddying this issue is the lack of a standard analytical method for determining the empty-to-full capsid ratio.

One of the most commonly used methods to quantify the empty-to-full ratio is analytical ultracentrifugation (AUC). However, the low throughput associated with AUC has decreased its usage [6], [7]. Alternatively, qPCR coupled with ELISA returns the vector genome (vg) concentration and capsid concentration, respectively. However, qPCR has been demonstrated to have high variability and interassay variation when used for AAV genome quantification over the past few years [8]–[10]. Droplet digital PCR (ddPCR) has recently emerged as a more robust alternative to qPCR, largely owing to its independence from a standard curve for analysis. ddPCR, however, necessitates the purchase of highly specialized droplet generators, thermalcyclers, and readers, which not only increases the cost but also requires a long turnaround time of up to 8 hours [11]. Extensively long time-to-result assays can become process development bottlenecks, hampering nimble, iterative process improvements.

Here, we demonstrate that CRISPR Diagnostics (CRISPR-Dx) could be a suitable approach to quantify AAV genome titers more robustly than qPCR and ddPCR, but requiring significantly less time and cost. CRISPR-Dx has recently emerged as a powerful tool for rapid, isothermal, and quantitative nucleic acid detection. Cas12a (Cpf1) [12] and Cas13 (C2c2) [13] are endonucleases of bacterial origin that exhibit ssDNA and RNA cleavage, respectively. The unique nonspecific nucleic acid *trans*-cleavage by Cas12/Cas13 is activated upon *cis*-cleavage of a target DNA sequence for Cas12 and RNA sequence for Cas13a complementary to its CRISPR RNA (crRNA). CRISPR-Dx has been extensively demonstrated as a powerful tool for pathogen detection [14], [15], typically at the point-of-care. For example, two diagnostic tests for SARS-CoV-2 were granted emergency use authorization by the FDA [16] during the COVID-19 pandemic. Herein, we describe the design and development of the CRISPR-AAV Evaluation (CRAAVE) assay for quantitative AAV vector genome titer measurement with different constructs.

We carefully constructed the CRAAVE assay to maximize sensitivity, minimize time-to-result, and provide a potentially universal design that can be used to quantify multiple transgene constructs encapsidated within multiple serotypes (e.g., AAV2, AAV5, and AAV6). AAV sample preparation begins with DNase treatment, a step that is needed only for unpurified samples (upstream samples), to digest nonencapsidated DNA containing the transgene sequence (**Figure 1**). Next, the transgene will be exposed for detection, accomplished by lysing viral capsids with a short, high temperature incubation at 95°C for 15 mins. The viral lysate is then assayed with the CRAAVE assay employing Cas12a as the effector. Upon Cas12a specific binding and subsequent cleavage of the AAV genome, Cas12a begins nonspecifically cleaving ssDNA fluorophore and quencher (F-Q) reporter molecules, generating fluorescence signals that are proportional to the concentration of AAV genome (**Figure 1**). The optimized CRAAVE assay can detect AAV genomic titers as low as 7e7 vg/mL, has a working range over 1 log of genome titers, and only requires 30 minutes of reaction time. The assay can be used for quantitation of both upstream and downstream AAV samples. The assay demonstrated a high analytical accuracy (< 3% error) when quantifying unknown AAV titers with a pre-established calibration curve constructed from AAV containing the same genetic construct.

**Figure 1:**
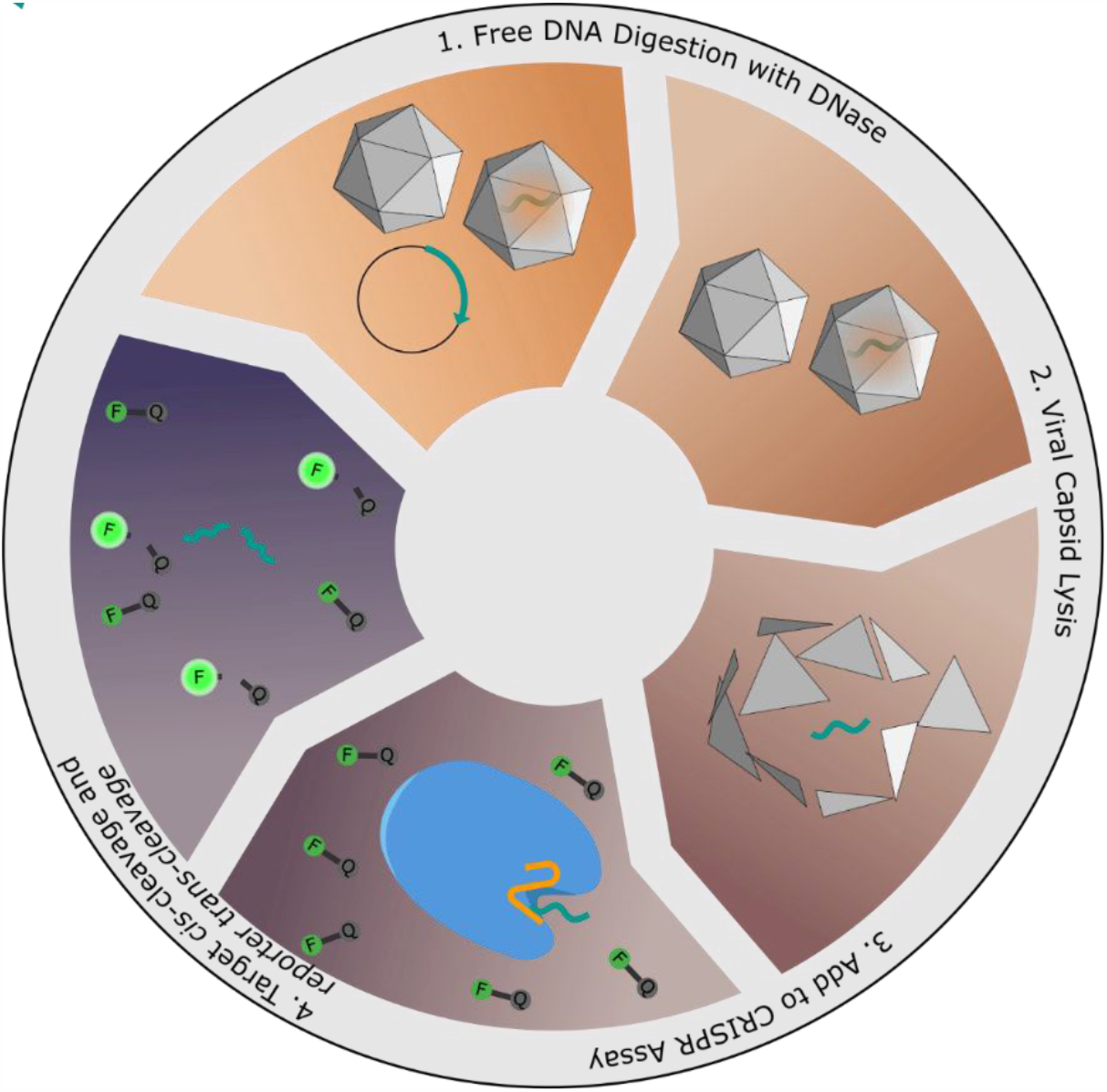
Schematic of CRAAVE Assay Workflow. F-Q: fluorescence-quencher.

## Results

rAAV constructs share commonality through obligatory inclusion of endogenous AAV genetic sequences and inverted terminal repeats (ITRs) at distal ends of the recombinant genome (**Figure 2a**, red and blue sequences). The highly conserved AAV type 2 (AAV2) ITR sequences are 145 nucleotide long hairpin structures required for replication as well as genome packaging [17]. AAV2 ITR sequences are the most widely studied and characterized ITRs among all serotypes. Thus, AAV2 ITRs are predominately used in rAAV constructs. The D-sequence, an element of the ITR that does not contribute to the hairpin structure, is a 20 nucleotide long sequence, which serves as the signal for capsid packaging and replication [18] (**Figure 2a**, blue sequence). From a genome optimization perspective, the D-sequences should remain in their wild type to lead to high levels of genome encapsidation. We designed our CRAAVE assay to be specific to the AAV2 ITR D-sequences to eliminate the requirement for crRNA redesign for various transgenes (**Figure 2a**). Additionally, we avoided coupling our assay to any preamplification steps, a common strategy taken to boost the sensitivity of CRISPR assays [14], [15], to maintain minimal time and complexity required for running the assay.

**Figure 2:**
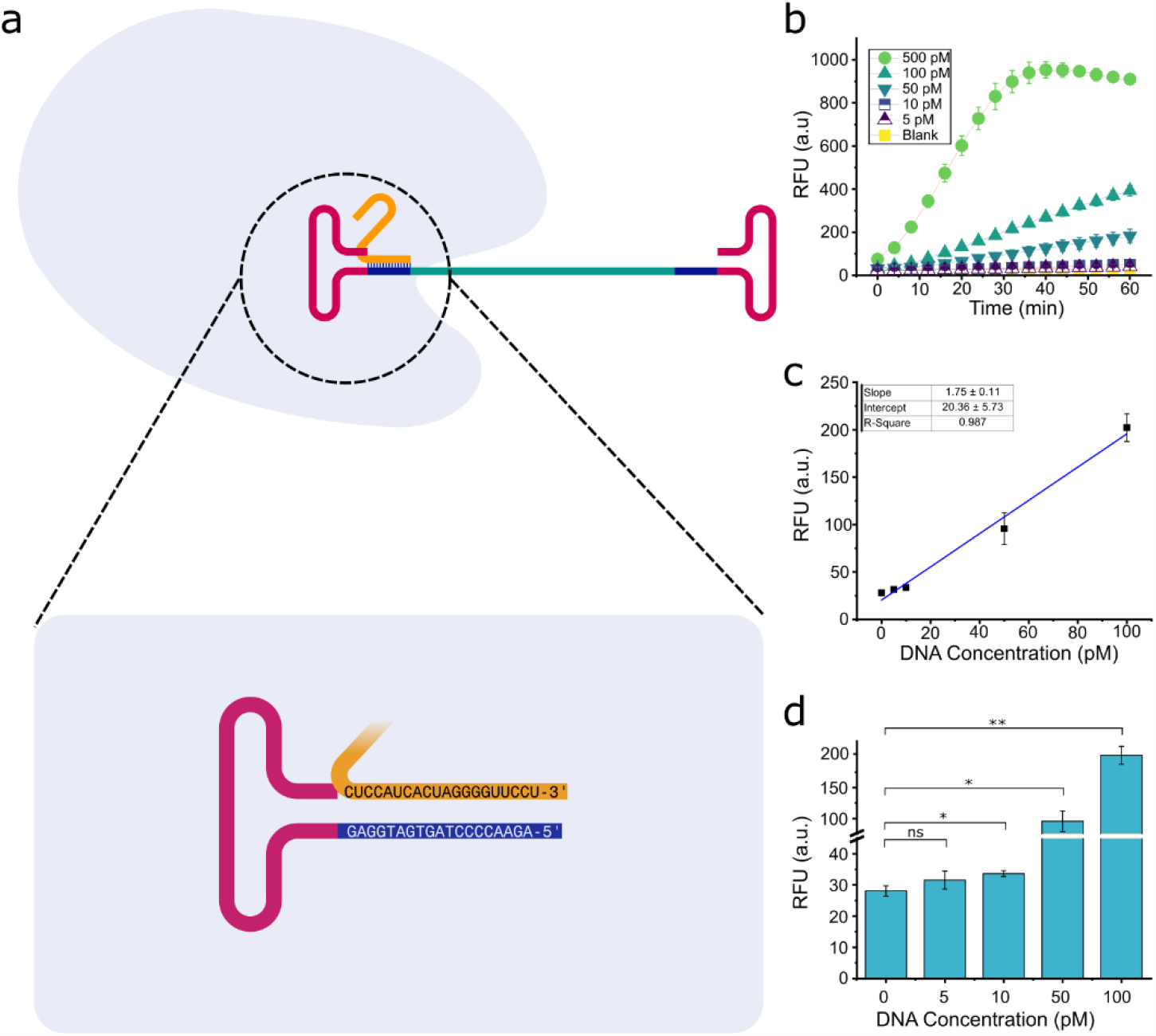
Assay Design and Proof of Concept. (a) The CRAAVE assay is constructed to specifically detect the AAV2 ITR D-sequence (navy blue region). After binding and cis-cleavage of the D-sequence, the activated Cas12a enzyme begins nonspecific degradation of ssDNA reporter molecules. (b) Real-time CRISPR assay results on a 42 nt synthetic AAV D-sequence target, where from (c) a fitted linear calibration curve after 30 minutes of reaction, a limit of detection (LOD) of 8 pM was achieved. (d) Assay intensities after 30 minutes reaction in a plate reader where error bars represent standard deviation (n = 3). Statistical analysis was performed using a two tailed student’s t-test, where ns = not significant with p > 0.05, and asterisks (* = p < 0.05, ** = p < 0.005) represent significant differences.

### Proof of Concept and crRNA Design

We utilized LbCas12a, one of the commonly used Cas12a orthologs, because it has a well-demonstrated high rate of signal generating *trans*-cleavage [19], [20], the mechanism causing signal transduction for the CRISPR assay. The assay reporter molecule is a dual labelled short 5 nt ssDNA with a FAM fluorophore and complementary quencher at the distal ends. The crRNA for LbCas12a is 40-44 nt in length, with a conserved repeat region of 21 nt, and a customizable spacer region of 20 – 24 nt [21]. We designed a 20 nt spacer sequence **(Table S1)** specific to the single stranded D-sequence **(Figure 2a)** for potentially universal quantitative ability. Two D-sequences are present on each end of the AAV ssDNA genome. Our crRNA is specific to the D-sequence of the ITR in the “flip” orientation [22]. A similar universal strategy was demonstrated for qPCR [23]. However, qPCR amplicons are longer than 20 bp. As a result, amplification was performed within the hairpin region of ITR, leading to variable results due to secondary structure [9].

We first wanted to test the feasibility of our detection system on a short synthetic 42 nt ssDNA target containing the D-sequence as well as minor secondary structure mimicking the structural effect of the hairpin **(Table S1)**. We found that the CRAAVE assay provided fairly sensitive detection of the synthetic sequence, with a limit of detection (LOD) of <8 pM after 30 minutes of reaction **(Figure 2b,c)**. LODs for preamplification-free DNA detection approaches are typically in the picomolar level after 1-2 hours of reaction according to the literature [14], [20], [24]. Thus, we were surprised to see a high level of sensitivity within half an hour. The proof of concept demonstrates that a CRISPR diagnostics approach provides impressive sensitivity on synthetic AAV target as well as the robust performance of engineered detection components.

Next, we wanted to determine assay functionality on real AAV samples. We obtained AAV9-CMV-GFP from a commercial vendor, Virovek. A simple pretreatment step similar to those reported in AAV PCR method development studies was adopted [8], [10], [25]. Briefly, DNase I was added to a final concentration of 0.5 U/μL reaction and incubated at 37°C for 30 minutes. Then, the viral capsids were lysed at 95°C for 15 minutes (**Figure 1)**. We chose not to use Proteinase K not only because heating at 95°C lyses capsids efficiently, but Proteinase K added a significant amount of error to the assay results **(Figure S1)**. Next, we performed multiple crRNA candidate screenings for maximized AAV detection sensitivity. In total 215 unique crRNA candidates within the ITR sequences were computationally screened for binding efficiency to the AAV genome using the NUPACK 4.0 analysis software **(Figure S2a**,**b)**. crRNA sequences were generated via a combinatorial approach by assessing candidates between 20 and 24 nt long and systematically adjusted inset (0-22 nts) by increasing distances into the ITR loop region **(Figure S2a**,**b)**. Candidate strength for AAV ITR binding was assessed via predicted percentage crRNA and AAV genome binding (**Figure S2c**). A secondary experimental crRNA screening to challenge the NUPACK 4.0 candidate predictions was performed on 6 selected promising candidates, including the flip ITR crRNA (**Figure S2d**). Interestingly, the flip ITR crRNA was the best experimental performer **(Figure S2e)** despite being predicted to be the least prolific AAV genome binder out of the 6 crRNAs in the computational screening. We attributed the minor discrepancy between the computational prediction and experimental validation to the imperfect nature of the current simulation package, which is mainly designed to study DNA-DNA interaction instead of CRISPR/Cas guided DNA-RNA binding in this work.

Using the selected flip ITR crRNA sequence, we then challenged the CRAAVE assay against Virovek AAV samples serially diluted down to 1e8 vg/mL **(Figure 3a)**. The assay displayed a linear relationship between 1e8 and 1e10 vg/mL with strong linearity (R^2^ = 0.977) **(Figure S3a)**. We were surprised by the difference in sensitivity of the CRISPR assay to real AAV as opposed to a short synthetic sequence. Based on the titers reporter by Virovek, the LOD of real AAV genome was calculated to be 8.4e7 vg/mL (or 140 fM), even better than the previous synthetic target (LODs of 8 pM, **Figure 2 c&d**), over an order of magnitude increase in sensitivity. Genome concentrations higher than 1e10 vg/mL exhibited nonlinearity, possibly due to the hook effect of high analyte concentration **(Figure S3b)**. A few earlier studies suggested that some chemical additives such as L-proline and Betaine can help enhance CRISPR diagnostics performance, although the mechanism is not well understood [26], [27]. We tested these chemicals along with urea as a potential method to alleviate genome secondary structure to extend the working range beyond 1e10 vg/mL. While the chemical treatments increased linearity slightly, there wasn’t a significant benefit to sensitivity from the additive chemicals tested **(Figure S3c)**. Although Virovek vectors are purified, we noticed a significant difference in fluorescence when treated with and without DNase **(Figure 3b)**, suggesting the presence of free AAV transgene present outside the capsids, either in the form of transfer plasmid DNA or significant unintended release of vector genomes. When treated with 1 U/μL DNase, 1e9 vg/mL AAV yielded an average signal fold change of 1.27, whereas no DNase added yielded a 1.82-fold change compared to the blank. 2e9 vg/mL after treatment with 1 U/μL DNase yields a signal fold change of 1.5, leading to a rough estimate that over two-fold more sequences containing the ITR are present outside the capsids in Virovek’s product. The DNase treatment was challenged against 3.6x10^12^ copies/mL transfer plasmid **(Figure 3c)**. Based on fluorescence change from no template control to no DNase added, 0.1 U, 0.5 U, and 1 U DNase/μL treatment reaction removed 81%, 84%, and 85% of the plasmid, respectively. The DNase step is commonly used as a cleanup step for any free DNA not already digested by Benzonase, commonly added to remove genomic DNA even in upstream processing [28]. Thus, it would be unlikely such a high amount of free transgene DNA would be present in samples for genome titer testing. Based on the plasmid digestion and Virovek DNase development results, 0.5U/μL reaction appears to be a conservative approach for cleaning up any free DNA present in analytical samples. Overall, the CRAAVE assay performed robustly on Virovek produced AAV as well as in the presence of DNase I enzyme.

**Figure 3:**
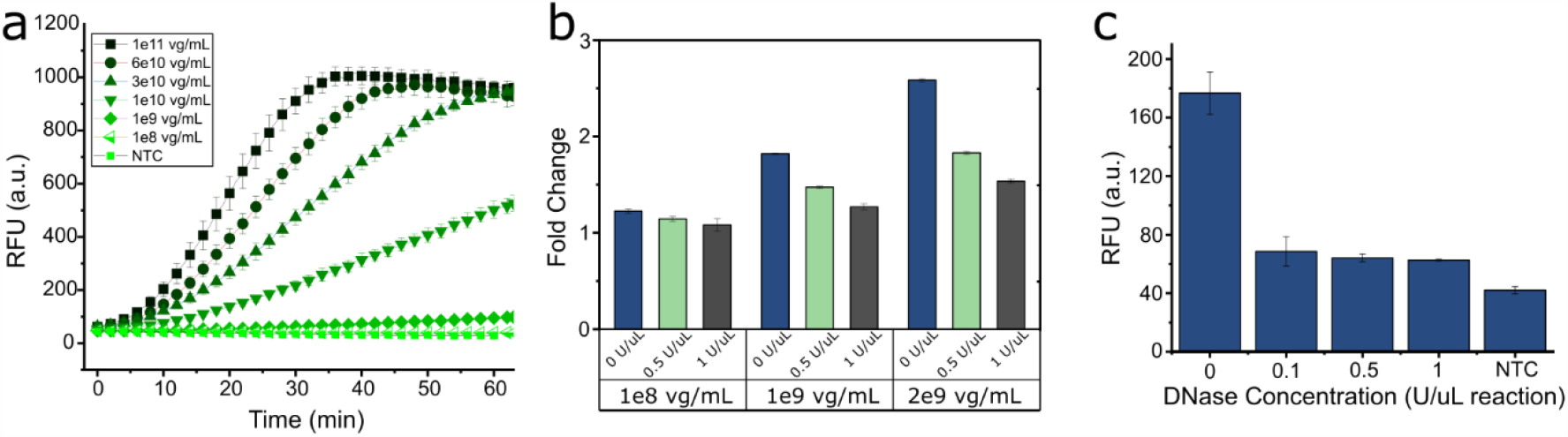
CRAAVE Assay demonstration on Virovek AAV. The CRISPR diagnostic assay for AAV was demonstrated on a commercially available AAV from Virovek prior to further assay development. (a) Real-time results of the CRISPR diagnostic assay on a Virovek AAV serial dilution containing a CMV-GFP construct. (b) Fluorescent signal intensity fold change after 30 minutes reaction time with 0, 0.5, and 1 U DNase per microliter of AAV pretreatment reaction for 3 different concentrations of AAV. Although Virovek manufactured AAV is reported to be purified, there was significant levels of DNA detected that was nonencapsidated. (c) DNase challenge against 3.6x10^12^ copies/mL of AAV transfer plasmid. DNase digested up to 85% (∼3x10^12^ copies/mL) of the plasmid.

### Assay Optimization

The CRAAVE assay was then optimized and tested on other AAV2 samples, such as affinity purified AAV2 produced using a HEK293 cell line triple transfection process, the most common technique for AAV production [29], [30]. Most CRISPR diagnostic assays are incubated at 37 °C. However, LbCas12a is known to be active at temperatures of up to about 50 °C. The CRAAVE assay was run against AAV2 targets at a concentration of 1e9 vg/mL at temperatures ranging from 30 °C to 65 °C to determine temperature effects on the assay **(Figure 4)**. With the exception of 65 °C, *trans*-cleavage was observed at all temperatures tested **(Figure 4a)**. The enzymatic reaction rate, interpreted from the slope of fluorescence versus time, increases up to 49 °C. But LbCas12a begins denaturing above this temperature, evidenced by decreasing slope at higher temperatures as well as no observable fluorescence at 65 °C. While thermophilic Cas variants have demonstrated *trans-*cleavage at temperatures of 60 °C and above coupled to LAMP pre-amplification [31]–[33], LbCas12a cannot typically retain activity at temperatures exceeding 50 °C. We also noticed there was a temperature dependence on the time required for *trans*-cleavage to commence **(Figure 4b)**. Fluorescence resulting from *trans*-cleavage was distinguishable from the control after 45 minutes, 11 minutes, and at the first read when the CRAAVE assay was run at 30 °C, 37 °C, and all higher temperatures other than 65 °C, respectively. We suspect this relationship is due to the relaxed secondary structure of the AAV DNA at higher temperatures (**Figure 4c,d**). Several reports of CRISPR detection of large, complex nucleic acid targets with secondary structure show a temporal delay with respect to observed fluorescence in real time kinetic curves of CRISPR detection [34], [35], a delay not typically observed when short nucleic acids are the CRISPR/Cas *cis*-cleavage target (**Figure S4**). This behavior could be attributed to a reduced efficiency in nucleic acid interrogation by the Cas enzyme as it needs to negotiate with significant secondary DNA structure. Since the assay kinetics benefit from increased temperature due to increased enzymatic reaction rate as well as reduced DNA structure leading to faster crRNA/DNA binding, we selected 45 °C as the temperature for further assay development.

**Figure 4:**
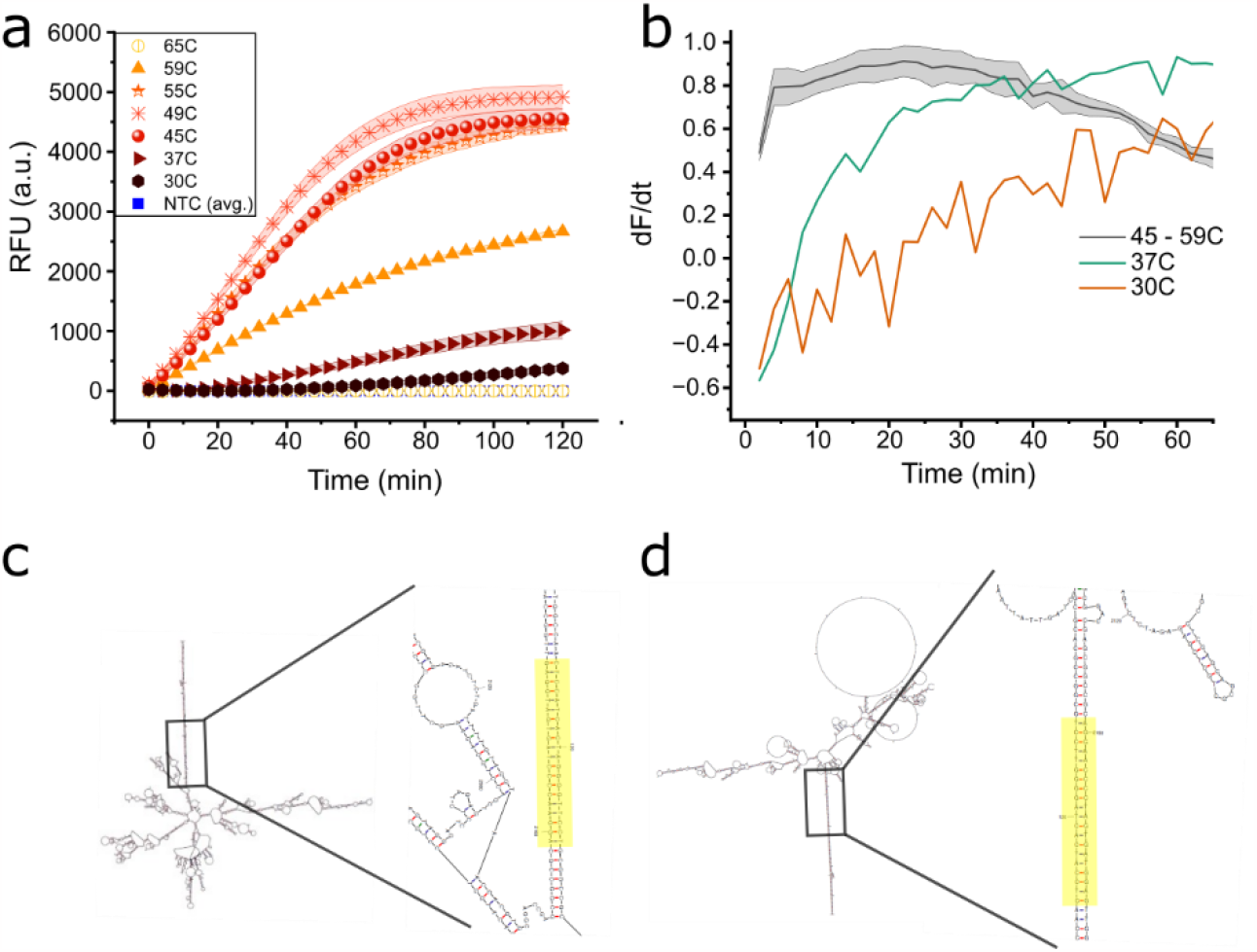
Effect of assay temperature on detection kinetics. The CRAAVE assay was developed by running at various temperatures with 1e9 vg/mL AAV genomic titer. (a) Real-time kinetic results show behavioral difference between the assay when run at 30 and 37 °C versus 45, 49, 55, and 59°C. Cas12a reporter cleavage for lower temperatures displays a temporal delay, significantly lowering detection sensitivity for a short assay time scale. (b) Visualization of normalized slope of fluorescence curves for all temperatures, with the exception of 65°C. Temperatures at 45°C and above reach near maximal slope within 5 minutes of assay initialization, whereas lower temperature normalized slopes do not reach maximal slope within 1 hour, a behavior that is likely due to the secondary structure of AAV genome. (c,d) Mfold-predicted AAV genome structures at (c) 37°C and (d) 45°C with the D-sequence crRNA binding site highlighted in yellow.

With the assay temperature selected at 45°C, we next wanted to determine the optimal concentration of the Cas ribonucleoprotein (RNP) for sensitive AAV quantification. We observed that a lower concentration of Cas RNP yielded higher rates of *trans*-cleavage at lower vector genome concentrations, but conversely, a higher concentration of Cas RNP yielded higher rates of *trans*-cleavage at higher vector genome concentrations **(Figure S5)**. Our aim was to develop an assay with the highest level of sensitivity for flexibility in quantitating different AAV sample types including lower titer upstream samples, so we focused on lower RNP concentrations for CRAAVE assay development. To this end, the affinity purified AAV2 samples were serially diluted to 4 concentrations between 1e8 vg/mL and 5e9 vg/mL and assayed with 1 nM, 2.5 nM, 5 nM, and 10 nM Cas RNP **(Figure 5a)**. Over the AAV genome titer range tested, there was a positive correlation between RNP concentration and sensitivity to AAV up to 5 nM RNP. 5 and 10 nM RNP provided very similar sensitivity within the AAV range tested. The positive correlation between target detection and Cas enzyme concentration has been intuitively reasoned that more available Cas enzymes for ssDNA reporter destruction lead to greater sensitivity [36]. Since 5 nM and 10 nM RNP exhibit similar sensitivities, we opted to proceed with 10 nM Cas12a RNP to provide a higher level of assay robustness in the event of unexpected Cas enzyme degradation during the assay.

**Figure 5:**
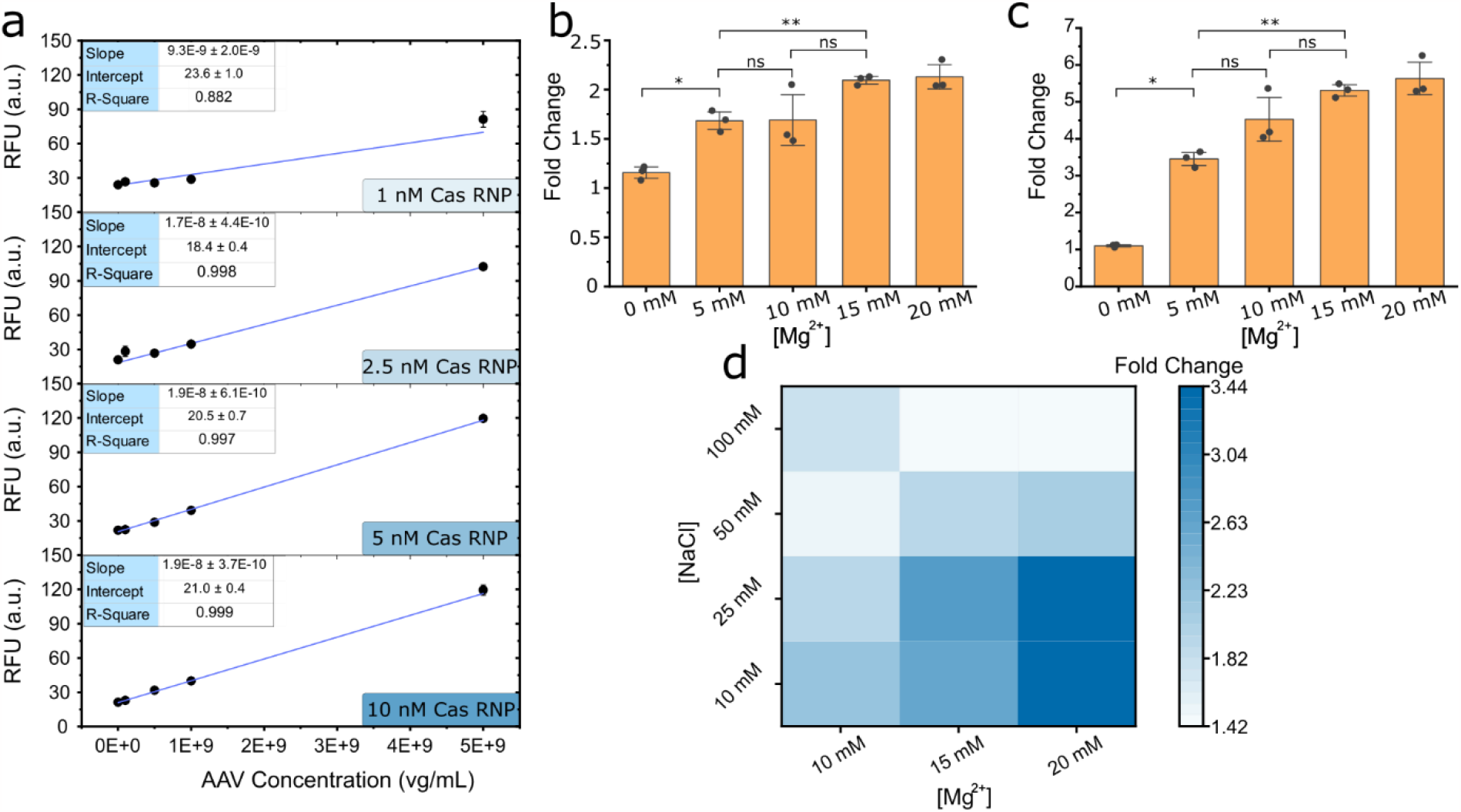
CRAAVE Assay Optimization on Affinity Purified AAV2: (a) The concentration of Cas enzyme and crRNA, together referred to as ribonucleoprotein (RNP) was tuned for most sensitive detection of AAV genomic titer. 4 RNP concentrations were tested against AAV genome titers ranging from 1e8 to 5e9 vg/mL. 10 nM RNP was selected was the most robust assay condition. Next, the standard NEBuffer 2.1, typically containing 10 mM Mg^2+^ was custom prepared to test the effects of modifying magnesium concentration. Results after (b) 30 minutes and (c) 60 minutes of reaction show that increasing the Magnesium above 10 mM provide additional sensitivity represented by signal fold change over the NTC, where error bars represent standard deviation (n = 3). Statistical analysis was performed using a two tailed student’s t-test, where ns = not significant with p > 0.05, and asterisks (* = p < 0.005, ** = p < 0.0005) represent significant differences. (d) Buffer composition effect on CRAAVE assay sensitivity was further assessed by modifying both NaCl and MgCl_2_ concentrations. Significantly better sensitivity was achieved when total ionic strength was lowered, but magnesium concentration was kept at 20 mM.

One of the most common reaction matrices for Cas12/13 diagnostic assays is NEBuffer 2.1 [20], [37]–[40]. We hypothesized this standard cleavage buffer could be further optimized. Mg^2+^ is one of the most important ions necessary for Cas12-a mediated cleavage and a concentration around 15 mM has been shown to be optimal for efficient *trans-*cleavage [19], [41], thus we suspected that an Mg^2+^ concentration higher than 10 mM included in standard 1x NEBuffer 2.1 would likely lead to faster ssDNA cleavage. Indeed, increasing the Mg^2+^ concentration up to 15 mM increases *trans*-cleavage rates **(Figure 5b)** significantly above the rate at 10 nM magnesium. Increasing the magnesium concentration above 15 mM to 20 mM did not significantly alter the *trans*-cleavage rate. We next explored the effect of ionic strength on the *trans*-cleavage rates more broadly by varying the magnesium and sodium chloride concentrations concurrently **(Figure 5c)**. Cas12a detection suffered at the highest ionic strengths tested (100 mM NaCl with Mg^2+^ at 15 and 20 mM), as the minimum fluorescence fold changes were observed. Interestingly, the maximum fold change was reached when Mg^2+^ was increased to 20 mM, but NaCl was decreased to 25 mM, suggesting that magnesium concentrations higher than 15 mM may be optimal for CRISPR diagnostics applications, but the total ionic strength may need to be decreased for this optimum to be realized. 25 mM NaCl and 20 mM MgCl_2_ provided significantly better detection results than all other conditions with the exception of 10 mM NaCl/20 mM MgCl_2_ and 25 mM NaCl/15 mM MgCl_2_, two conditions with comparable performance **(Figure S6)**. The optimized buffer condition of 10 mM Tris-HCl, 25 mM NaCl, and 20 mM MgCl_2_ was used for all subsequent CRAAVE assays conducted in microtiter plates.

### Assay Demonstration on Different AAV Serotypes and Quantification of Unknown Samples

The design of the crRNA is such that universal quantification of rAAV vectors utilizing AAV2 ITRs is theoretically possible. We challenged this hypothesis by running the CRAAVE assay against a serial dilution of 3 affinity purified AAV serotypes in a plate reader at 45°C. The genes of interest (GOIs) in AAV2, AAV5, and AAV6 were Cell Biolabs pAAV-GFP, Plasmid Factory pAAV-ssGFP, and a combination of both transgene plasmids, respectively. The CRAAVE assay exhibited slightly different sensitivities to each construct, serotype, or combination of both factors **(Figure 6)**. The calculated LODs were 7.1e7, 8.1e7, and 7.4e7 vg/mL for AAV2, AAV5, and AAV6, respectively (**Figure S7**). This represents an increase in sensitivity of ∼60 fold over the synthetic DNA target (**Figure 2b**). A portion of this enhancement can be attributed to assay optimization efforts. However, the majority of the increase is likely due to Cas enzymes’ affinity for larger DNA sequences, [37], [41], [43] an observed behavior for which the mechanism is not understood, but is under investigation.

**Figure 6:**
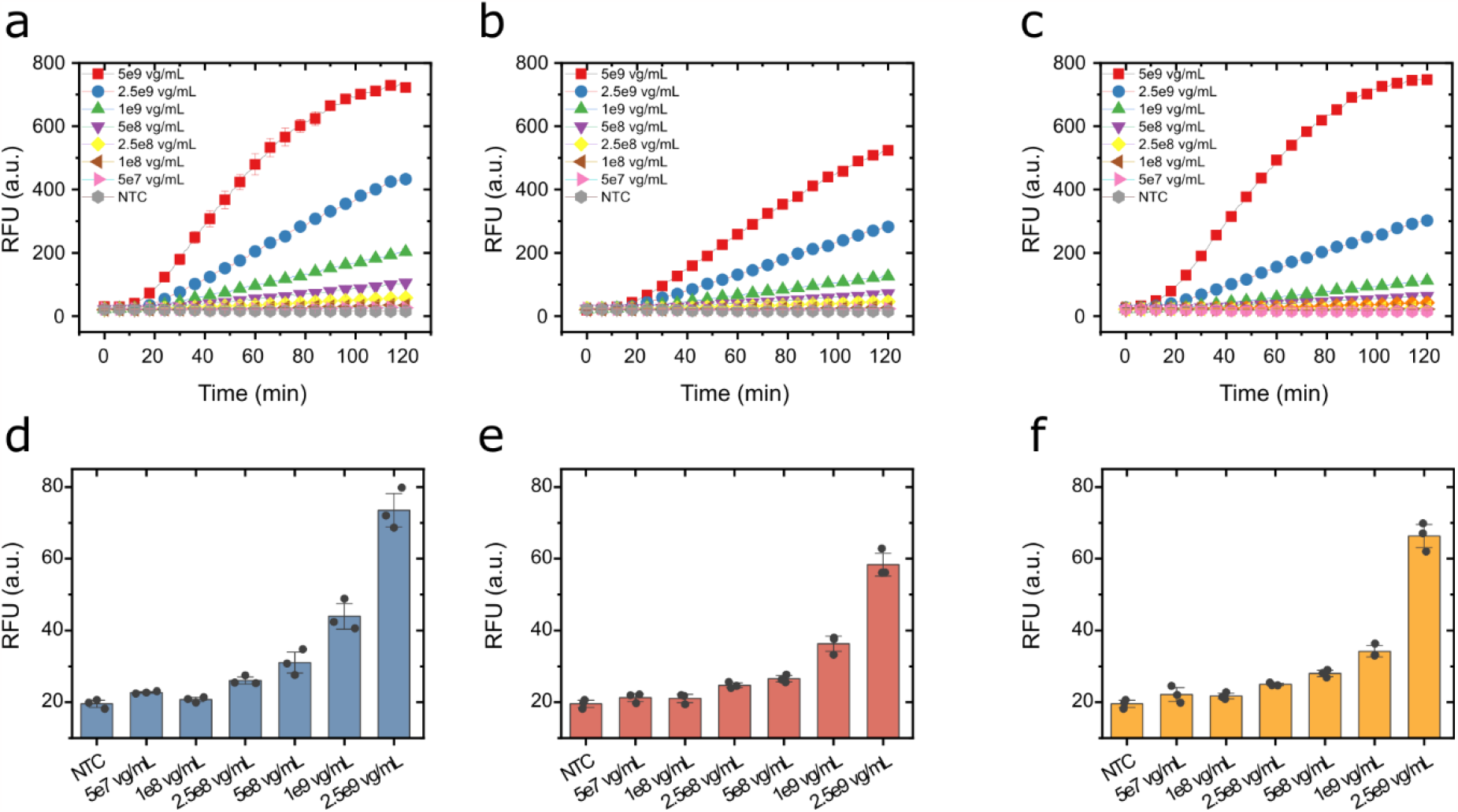
CRAAVE Assay Detection of different genomic constructs in 3 different affinity purified AAV serotypes. The developed CRISPR assay was challenged against (a,d) AAV2, (b,e) AAV5, and (c,f) AAV6, that encapsidated two different recombinant constructs in a microplate format read in a plate reader. The CRAAVE assay was able to generate signal against different constructs due to its specificity for the conserved ITR D-sequence.

To further investigate the large difference in kinetic rates observed for our real affinity purified AAV targets versus significantly shorter synthetic DNA, we assessed the catalytic rates via Michaelis-Menten kinetics (**Figure 7 and S8**). To this end, equimolar concentrations of synthetic target and rAAV2,

**Figure 7:**
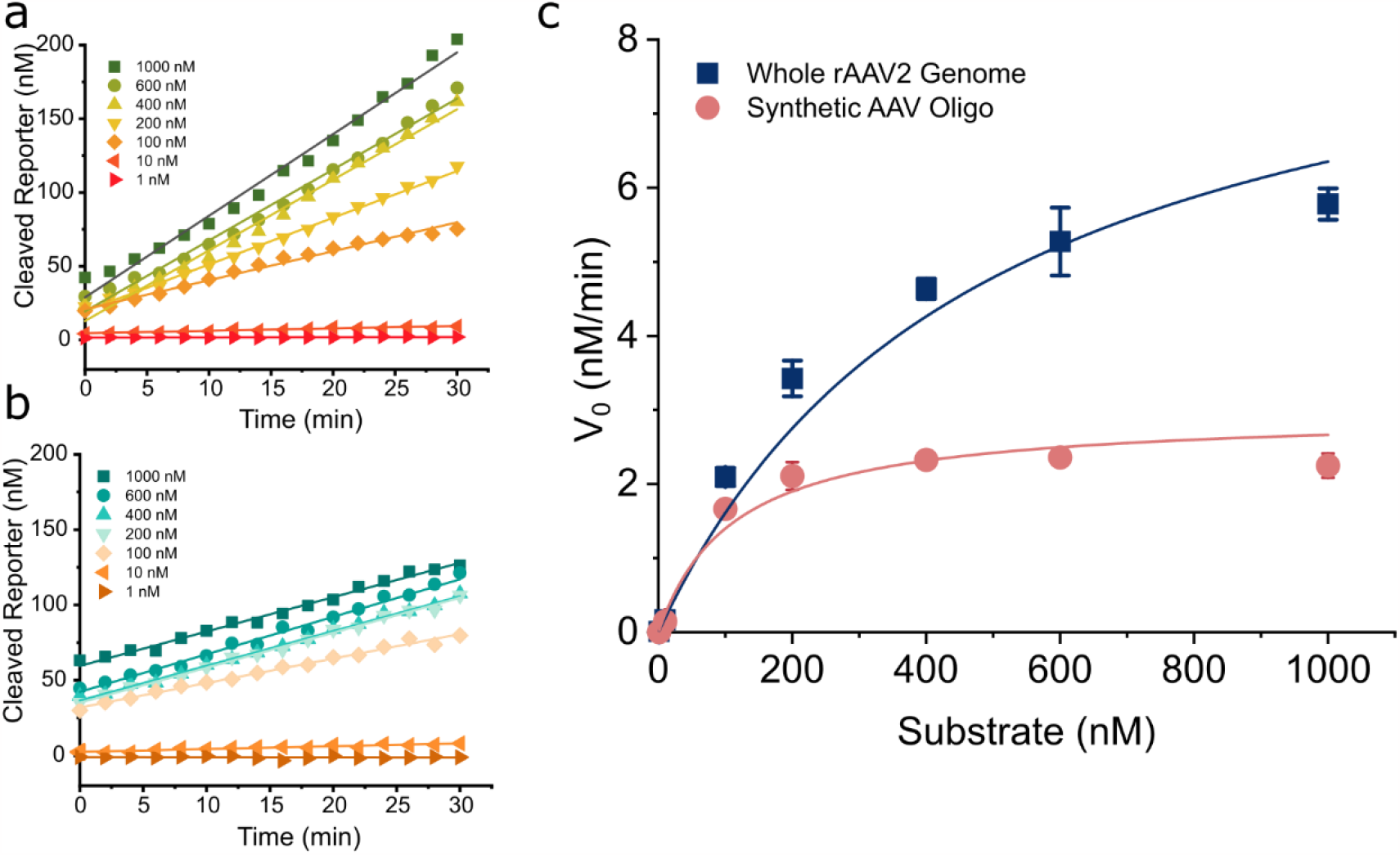
Michaelis-Menten Kinetics of the CRAAVE assay. Enzyme kinetic rates were investigated for Cas12a on a short synthetic sequence containing the ITR D-sequence and a much longer rAAV2 genome. (a) Representative reporter cleavage plot on AAV2 genome. (b) Representative reporter cleavage plot on synthetic AAV2 ITR sequence. (c) Overall Michaelis-Menten kinetics for both target types. Data were averaged from 3 replicates (n = 3). Enzymatic turnover rate was calculated to be 6.17 s^-1^ when activated with the synthetic oligo, and 19.67 s^-1^ when activated with the rAAV2 genome.

8.3 pM and 5e9 vg/mL, respectively, were mixed with the CRISPR assay with reporter concentrations ranging from 1 nM to 1000 nM. As observed in CRISPR detection of rAAV in the plate reader at 45°C, there is a temporal delay of roughly 15 minutes prior to observed AAV detection. Notably there is no delay at 45°C when performed in the thermal cycler **(Figure 4a)**. Therefore, initial velocities of AAV2 targets were assessed starting after 15 minutes **(Figure 7a)**. Initial velocities of synthetic AAV targets were assessed starting at time 0, since the reaction commences without delay **(Figure 7b)**. The average velocities from 3 replicates of each titrated reporter cleavage reaction were fitted nonlinearly **(Figure 7c)** to determine maximum velocity (V_max_) and Michalis-Menten constant (K_m_). Nonlinear regression was used to determine K_m_, which in turn was used to determine the turnover rate (k_cat_). The turnover rate (k_cat_), calculated as V_max_/[Enzyme], where enzyme concentration is total activated enzyme, limited by activating DNA concentration (8.3 pM) was determined to be 18.96 s^-1^ for AAV2 and 5.94 s^-1^ for the synthetic AAV oligo. The catalytic efficiency (k_cat_/K_m_) for AAV2 and the synthetic oligo was calculated to be 3.9e7 M^-1^s^-1^ and 5.1e8 M^-1^s^-1^, respectively. Complete kinetic values can be seen in **Figure S8**. Cas12a diagnostic assays sensitivities are kinetically limited by the catalytic turnover rate; the observed turnover rate when activated by AAV2 (whole genome) is nearly an order of magnitude higher than other broadly accepted k_cat_ reported literature values for amplification-free Cas12a diagnostics[12], [41], [44], [45].

We next demonstrated the quantitative power and flexibility of the CRAAVE assay by titrating a dilution of the AAV2 sample (**Table 1** and **Figure S9**) using the CRAAVE assay assembled in a standard microplate as well as after lyophilization on a chip-based format. The lyophilized chip format used wells holding up to ∼20 μL punched into polydimethylsiloxane (PDMS) bonded to glass slides. CRISPR reagents were added into each well, the entire chip lyophilized, then AAV samples added the next day. A photo of the actual chip can be seen in **Figure S10**. Fluorescence intensities of the chips were read on a benchtop fluorescence microscope (**Table 1** and **Figure S10b**). We benchmarked the CRAAVE assay against a standard qPCR method utilizing ddPCR titered AAV2 as a standard[10], [26]. From 3 dilutions using citrate buffer above the limit of detection (LOD) of the CRAAVE assay in both microplate and chip format, 2 out of 3 conditions accurately quantified the correct concentration of 7e9 vg/mL within 10% of the actual titer determined by ddPCR (**Table 1**). 5 out of the 6 qPCR dilutions quantified diluted AAV within the same error range. The 1:100 dilution for the CRAAVE assay fell outside of the limit of quantification (LOQ) for both assay formats. Thus its inaccuracy wasn’t considered. For both the CRAAVE and qPCR assays, AAV was added as one tenth the volume (1:10 dilution) of each reaction. Modifying the CRAAVE assay to accommodate a 1:5 dilution would allow more dilutions within the working range.

**Table 1:** CRAAVE Assay and qPCR quantitative comparison. Vg/mL in assay indicates the actual AAV titer in the assay condition as determined from ddPCR titration, and calculated titer represents the average titer (n=3) calculated from the standard curve multiplied by the dilution factor. %CV is represented as Std. Dv./mean*100.

| AAV Dilution Factor | vg/mL in assay | On-Chip CRAAVE Assay |  |  | Microplate CRAAVE Assay |  |  | qPCR Assay |  |  |
| --- | --- | --- | --- | --- | --- | --- | --- | --- | --- | --- |
|  |  | Calc. Titer | %CV | % Error | Calc. Titer | % CV | % Error | Calc. Titer | % CV | % Error |
| 1:10 | 7e8 | 6.81e9 | 6.2 | 2.7 | 7.22E9 | 3.00 | 3.2 |  |  |  |
| 1:20 | 3.5e8 | 6.46e9 | 4.6 | 7.7 | 6.00e9 | 3.16 | 14.3 |  |  |  |
| 1:50 | 1.4e8 | 3.24e9 | 6.3 | 53.7 | 7.03e9 | 0.72 | 0.4 | 7.23e9 | 0.28 | 3.2 |
| 1:100 | 7e7 | 1.22e9 | 10.6 | 83.6 | 3.97e9 | 1.14 | 43.4 | 7.37e9 | 0.39 | 5.3 |
| 1:1000 | 7e6 |  |  |  |  |  |  | 6.6e9 | 0.44 | 5.7 |
| 1:5000 | 1.4e6 |  |  |  |  |  |  | 6.89e9 | 0.17 | 1.6 |
| 1:10000 | 7e5 |  |  |  |  |  |  | 6.04e9 | 0.16 | 13.8 |
| 1:50000 | 1.4e5 |  |  |  |  |  |  | 6.78e9 | 0.19 | 3.1 |
| Working Range |  | ~1.5 logs |  |  | ~1.5 logs |  |  | ~8 logs |  |  |
| Time to Result |  | 30 minutes |  |  | 30 minutes |  |  | ~1 hour |  |  |
| LOD |  | 9.72e7 vg/mL |  |  | 6.97e7 vg/mL |  |  | 5e3 vg/mL[46] |  |  |
| LOQ |  | 3.72e8 vg/mL |  |  | 2.68e8 vg/mL |  |  |  |  |  |

## Discussion

Biomanufacturing process analytical technology (PAT) has seen an increased shift towards real-time monitoring since the FDA-encouraged increased adoption and focus on robust analytical technologies [17], [47]. AAV vector manufacturing has broadly not seen new analytical technologies co-develop with the emerging cell and gene therapy market. For these reasons, we developed a quantitative CRISPR assay for determining the total vector genome titer with a sensitivity suitable for upstream samples that could return a result in as little as 30 minutes.

All of the current rAAV genome quantification techniques have unique advantages as well as disadvantages, typically associated with sensitivity, long assay times, specificity, or sample volume required. qPCR emerged as a trusted analytical tool. However, reports of variation in reported titers soon followed, likely a result of variation between amplification efficiencies of plasmids for standardization and rAAV genomes. ddPCR is now regarded as superior to qPCR for accurate titration of rAAV genomes. The developed CRAAVE assay still requires a standard curve for interpretation of results (**Figure S7**). However, we propose using an rAAV reference as the standard, a technique successfully demonstrated for qPCR [10]. Given the universal nature of the CRAAVE assay, and differences in response to two genetic constructs tested, we would recommend that each construct be quantified with a standard-free method like ddPCR prior to application of the CRAAVE assay. Process development analytics would likely see the most benefit of the CRAAVE assay when optimizing process steps such as transfection, for example. The CRAAVE assay could return a result in as little as 30 minutes assay time when using a standard containing the same genetic construct. Additionally, the chip-based CRAAVE assay provides a highly flexible format for an off-the-shelf ready to use assay that requires no assay set-up, allowing end users to quickly titer rAAV samples with minimal time required for set-up and assaying.

The CRAAVE assay demonstrated limits of detection below 1e8 vg/mL in each case in 30 minutes of assay time. The CRAAVE assay also has superior flexibility in format and reader devices. The assay can be set up and read in a plate reader, thermal cycler, or any other device that can quantitate fluorescent intensities. As far as we are aware, CRISPR diagnostics has not yet been deployed as an analytical tool for biomanufacturing. The results of this study suggest that it could serve as a potential alternative to PCR methods for rapid quantitation of AAV titers.

## Methods

### CRISPR Diagnostics Assay Preparation and Execution

LbCas12a Ultra, all CRISPR RNAs and DNA oligos, and Fluorophore-quencher labeled reporter were purchased from Integrated DNA Technologies (IDT DNA). The single-stranded reporter DNA was constructed with a 5 nucleotide sequence, TTATT, 5’ labeled with 6-FAM and 3’ labeled with Iowa Black Quencher. Cas12a and crRNA working stocks were prepared by dilution to 5 μM in NEBuffer 2.1 and nuclease-free water, respectively, prior to storage at -80°C until needed for use. Cas RNP was first prepared by combining LbCas12a, crRNA, 10x NEBuffer 2.1, and nuclease-free water to achieve a 500 nM RNP concentration, which was subsequently incubated at room temperature for not less than 15 minutes. The prepared Cas RNP was then added to a master mix comprised of 10 nM Cas RNP, 400 nM FQ reporter, 1x NEBuffer 2.1 or optimized buffer, and nuclease-free water. The assay was run in a flat bottom, black 384-well plate in 30 μL reaction volumes in a SpectraMax M2 plate reader (Molecular Devices), quantitating FAM fluorescence using 490 nm excitation and 520 nm emission. Assay temperature kinetics study was performed in PCR tubes at 20 μL volume with the same assay concentrations as discussed above in a CFX Connect Real-Time System (Bio-Rad).

### qPCR AAV Quantification

qPCR quantification of AAV2 vg titer was accomplished with a primer/probe set targeting the GFP gene. Primers were purchased from Integrated DNA Technologies, and the Taqman-tamra hydrolysis probe was purchased from Fisher Scientific. Full sequence information can be found in **Supplementary Table 1**. Reactions were assembled at 25 μL volumes with Fast Advanced Master Mix (Fisher Scientific) and primer/probe concentrations of 200 nM. Thermal cycling and real-time detection were carried out in a CFX Connect Real-Time System (Bio-Rad). Thermal cycling conditions were 50°C for 2 minutes (UNG incubation), 95°C for 20 seconds (polymerase activation), followed by 40 cycles of denaturation at 95°C for 3 seconds and combined annealing/extensions steps at 60°C for 30 seconds.

### Chip Fabrication and Lyophilization

PDMS (Sylgard 184, Dow-Corning) was produced by mixing the base and curing agents at a 10:1 (w/w) ratio, and degassed by placing at 4°C for 15 minutes prior to curing at 65°C for ≥4 hours. The cured PDMS was then cut to roughly 25 mm in width and 60 mm in length. A 3 mm biopsy punch (MedBlades) was used to punch 40 wells through the PDMS using a 3D printed jig. The prepared PDMS was then bonded to a glass slide (Fisher Scientific) using a plasma cleaner (Harrick, PDC-32G) using 540 Joules of energy.

The CRISPR diagnostic master mix was prepared at a 5x concentration for lyophilization with additional magnesium included as well as 5% trehalose for stability during lyophilization. 2 μL of the lyophilization CRAAVE assay mix was added to each well, then frozen at -80°C. The chip was then placed in the lyophilizer (Labconco) for lyophilization at 0.2 mbar for several hours. The lyophilized chips can be used immediately or stored for later use in a dessicated sealed bag.

### Chip Assay and Readout

Chip assay execution was performed by diluting prepared AAV samples tenfold into the optimized CRISPR diagnostics buffer. The chip’s wells are segregated into a standard curve zone and sample titration zone with each concentration having 4 wells. 10 μL of each standard and sample for titration were added to each well with mixing, then sealed with 5 μL mineral oil. The chips were then incubated for 30 minutes at 45°C in a benchtop incubator.

The chips were read on a benchtop microscope (Olympus, BX43) using an LED excitation source with a GFP filter set consisting of an excitation filter with a 470 nm center wavelength (CWL) and 40 nm bandwidth & emission filter with a 525 nm CWL and 50 nm bandwidth. Imaging was performed with 250 ms exposure time and 4x objective. Intensities of each image were analyzed using ImageJ.

### Limit of Detection Analysis

Limits of Detection (LOD) were calculated using the following equation:

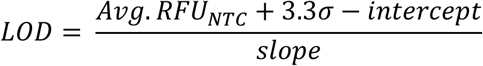

where the detection limit is analytically determined by calibration curve slope, intercept, and detector noise in the absence of target. Limits of quantification (LOQ) were determined using the same equation with the exception of the constant multiplying standard deviation (s). LOQ is assessed by multiplying s by 10 rather than 3.3.

### Michaelis-Menten Kinetic Analysis

Cas12a Michaelis-Menten kinetics experiment was carried out in a flat bottom, black 384-well plate in 30 μL reaction volumes read with the same instrument and settings as other CRISPR diagnostic assays. Reaction velocities for AAV2 genome were plotted with time = 0 starting after 15 minutes of the assay once detection began. The Michaelis-Menten plot was constructed by averaging reporter cleavage velocity replicates (n=3) generated for each point. Nonlinear fitting was performed in Origin 2023 to the Michaelis-Menten function, from which the V_max_ and k_m_ constants were extracted.

### AAV Production and pretreatment

Affinity purified AAV was pretreated by capsid lysis at 95°C for 15 minutes in a standard thermal cycler. AAV dilutions prior to CRISPR and qPCR titration were performed in 1x DNase buffer (Invitrogen) with

0.001% Pluronic F-68 or citrate buffer where noted. Citrate dilution buffer was comprised of 20 mM Citrate, 200 mM NaCl, 34 mM Bis-Tris Propane, pH 7, and 0.001% Pluronic F-68.

### AAV Purification

AAV crude harvests were centrifuged and filtered (0.22 μm) to remove residual cell debris. After clarification, each AAV was affinity chromatography purified using an ÄKTA Explorer 100 system. All buffers used during chromatography purification contained 0.001% Pluronic F-68. AAV2 was loaded onto Capto™> AVB (Cytiva) affinity resin and eluted with 20 mM sodium citrate, 200 mM NaCl, pH 2.5. Elution fractions were neutralized to pH 7 with 550 mM Bis-Tris propane, pH 10. AAV 5 and 6 were loaded onto CaptureSelect™> AAVX affinity resin. AAV5 was eluted with 100 mM phosphoric acid, pH 2.3 and neutralized with Bis-Tris propane as described above. AAV6 was eluted with 200 mM glycine, pH 2.0 and also neutralized with Bis-Tris propane.

## Data Availability

The data presented in this study, are included within the article and supplementary information. Further inquiry for original data, upon reasonable request, can be directed to the corresponding author.

## Acknowledgments

The authors would like to thank Merck KGaA and Millipore-Sigma for their support and supply of rAAV. The authors would also like to acknowledge the funding support from the National Institute for Innovation in Manufacturing Biopharmaceuticals (NIIMBL). We would also like to thank Christopher Cummings and Shriarjun Shastry from the Biomanufacturing, Training, and Education Center (BTEC) at NC State for their work in AAV production and purification to support this work.

## Author Contributions

ZH and QW conceived the project and designed the experiments. SM and AH provided input and discussion towards experimental design and manuscript revision. ZH, NL, NM, JK, LT, and DF performed the experiments. ALN, LO, HM, HG, and OR produced and purified the AAV. ZH and QW wrote the original manuscript, and all authors reviewed and edited the manuscript.

