## Supporting information for "Rapid Adeno-Associated Virus Genome Quantification with Amplification-Free CRISPR-Cas12a"

**Table S1:** crRNAs and sequences used in this study. Sequences are listed from 5' to 3'. The spacer region of the crRNAs is listed in bold.

| crRNA name | Sequence |
|---|---|
| Flip ITR crRNA | UAAUUUCUACUAAGUGUAGAU**CUCCAUCACUAGGGGUUCCU** |
| Flop ITR crRNA | UAAUUUCUACUAAGUGUAGAU**AGGAACCCCUAGUGAUGGAG** |
| Flip 20 nt inset 19 crRNA | UAAUUUCUACUAAGUGUAGAU**GCAGAGAGGGAGUGGCCAAC** |
| Flop 24 nt inset 18 crRNA | UAAUUUCUACUAAGUGUAGAU**AGUUGGCCACUCCCUCUCUGCGCG** |
| Flop 24 nt inset 15 crRNA | UAAUUUCUACUAAGUGUAGAU**UGGAGUUGGCCACUCCCUCUCUGC** |
| Flip 24 nt inset 18 crRNA | UAAUUUCUACUAAGUGUAGAU**CAGAGAGGGAGUGGCCAACUCCAU** |
| **Oligo name** | **Sequence** |
| GFP Fwd Primer | AGCAAAGACCCCAACGAGAA |
| GFP Rev Primer | GGCGGCGGTCACGAA |
| GFP Probe | CGCGATCACATGGTCCTGCTGG |
| FQ Reporter | /56-FAM/TTATT/3IABkFQ/ |
| Synthetic ITR ssDNA | AGGAACCCCTAGTGATGGAGTTGGCCACTTTTTAGTGGCCAA |

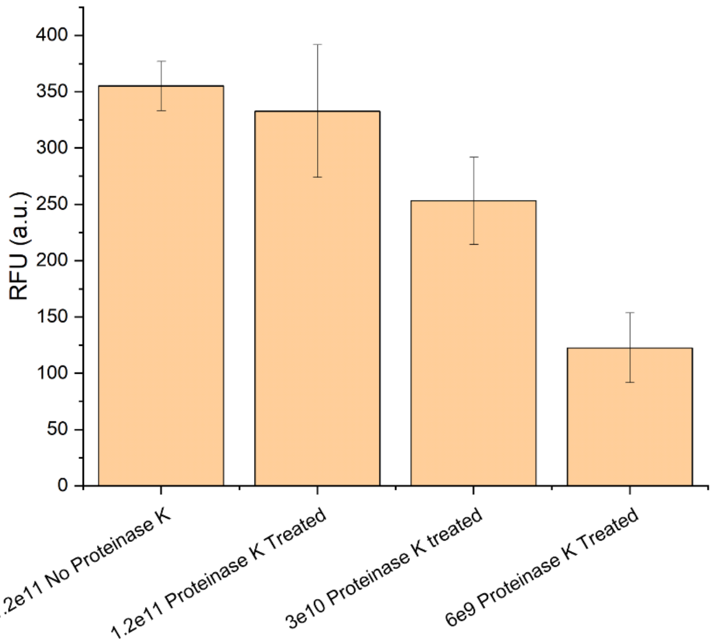


**Figure S1: Effects of Proteinase K treatment on CRISPR Assay.** Proteinase K when added as a capsid lysis agent significantly increased error in the CRISPR assay. Proteinase K treatment was removed, and capsids were lysed by heating at 95 $^{o}$C for 15 minutes.

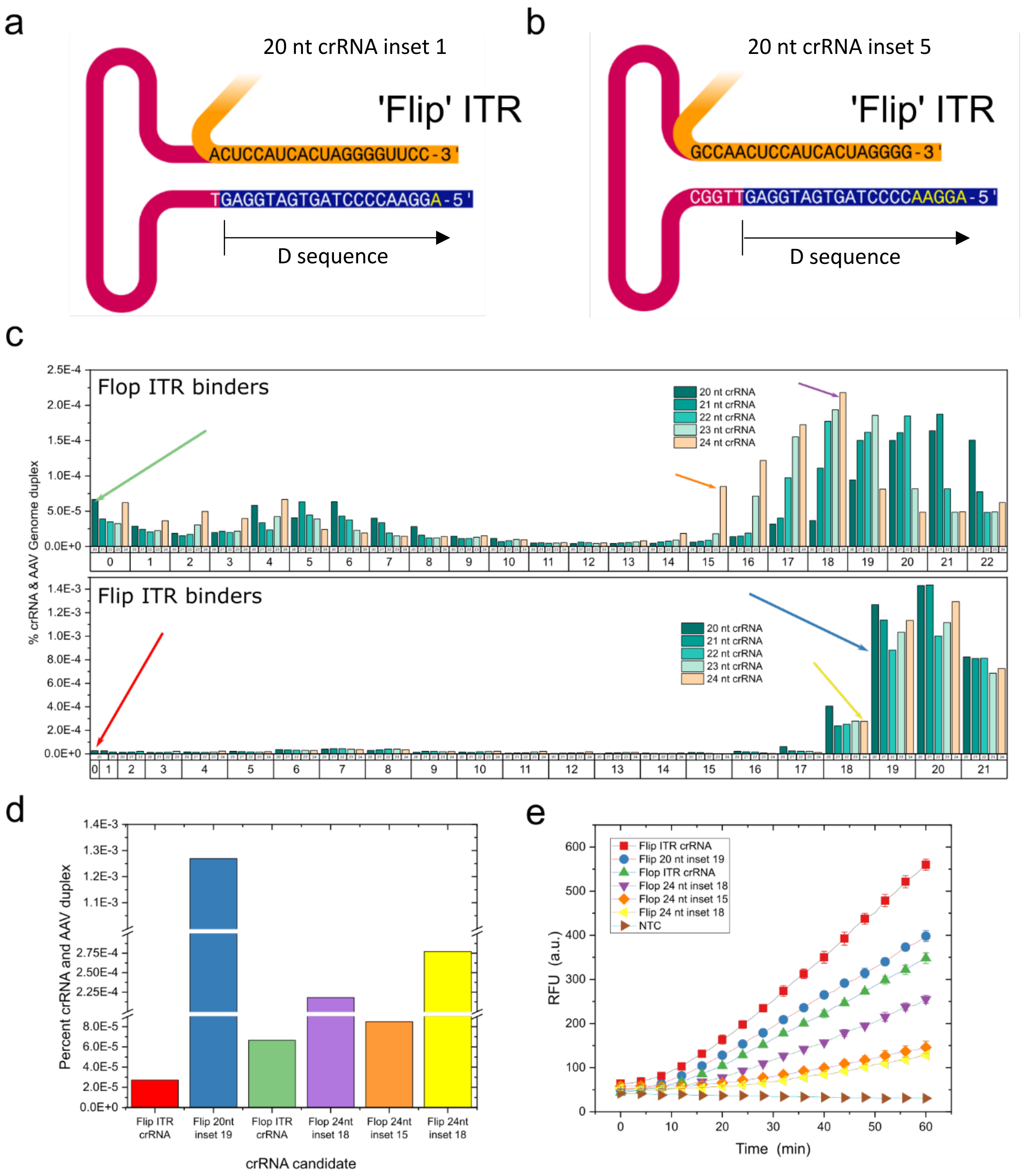


**Figure S2: NUPACK and experimental screening of AAV crRNA candidates.** 215 crRNA candidates were computationally screened using the NUPACK 4.0 package to predict free energy states of crRNA combined with rAAV genome. (a,b) Schematics depicting a 20 nt crRNA candidate with inset 1 and 5 nts into the ITR loop region. Note distance to the hairpin is not to scale. (c) Free energy, represented as percentage of crRNA candidates bound to the flip or flop orientation of the rAAV ITR. Candidates were generated by crRNAs between 20-24 nt long and with various inset distances into the ITR. (d) Top crRNA candidates selected from the computational screening process. (e) Experimental screening results of the 6 selected crRNA candidates. The 20 nt flip ITR crRNA binding directly to the flip orientation D-sequence outperformed all other candidates, despite being predicted to be least likely to bind.

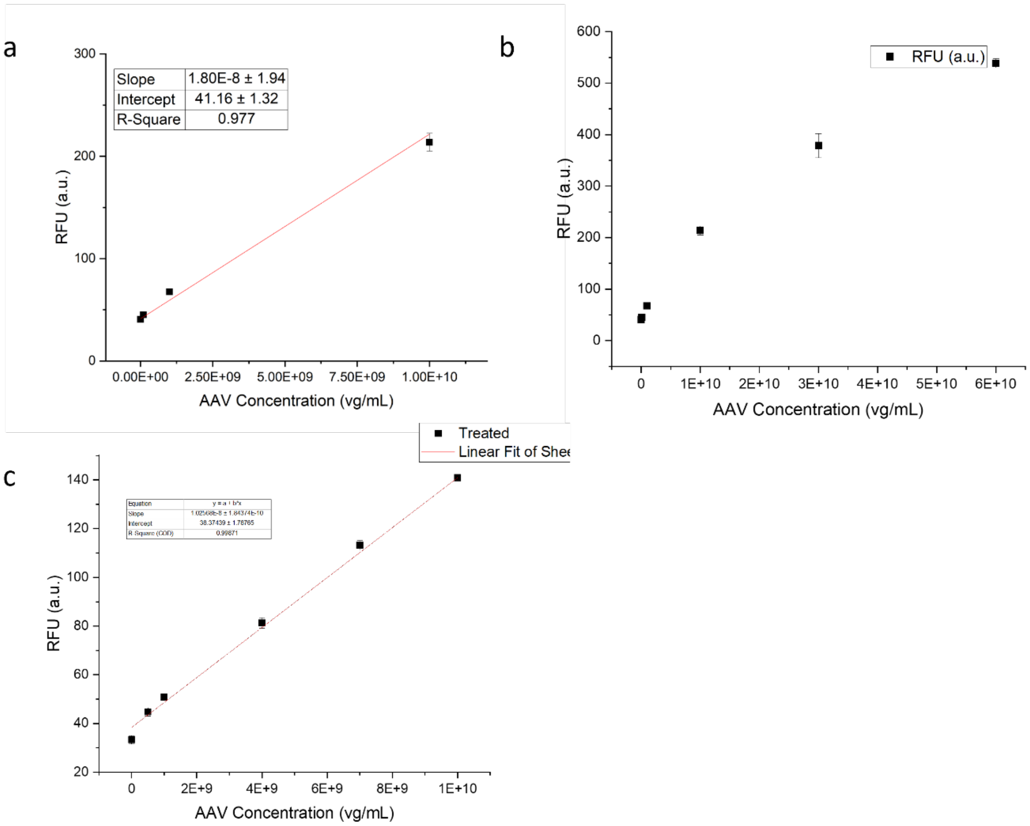


**Figure S3: CRAAVE Assay on Virovek AAV.** (a) Standard curve for Virovek AAV after 30 minutes assay time. Assay was run at 37 $^{o}$C with 30 nM Cas RNP. (b) Expanded standard curve exhibiting nonlinearity at concentrations above 1e10 vg/mL. (c) standard curve when the CRISPR assay included 500 mM Urea and 500 mM L-Proline.

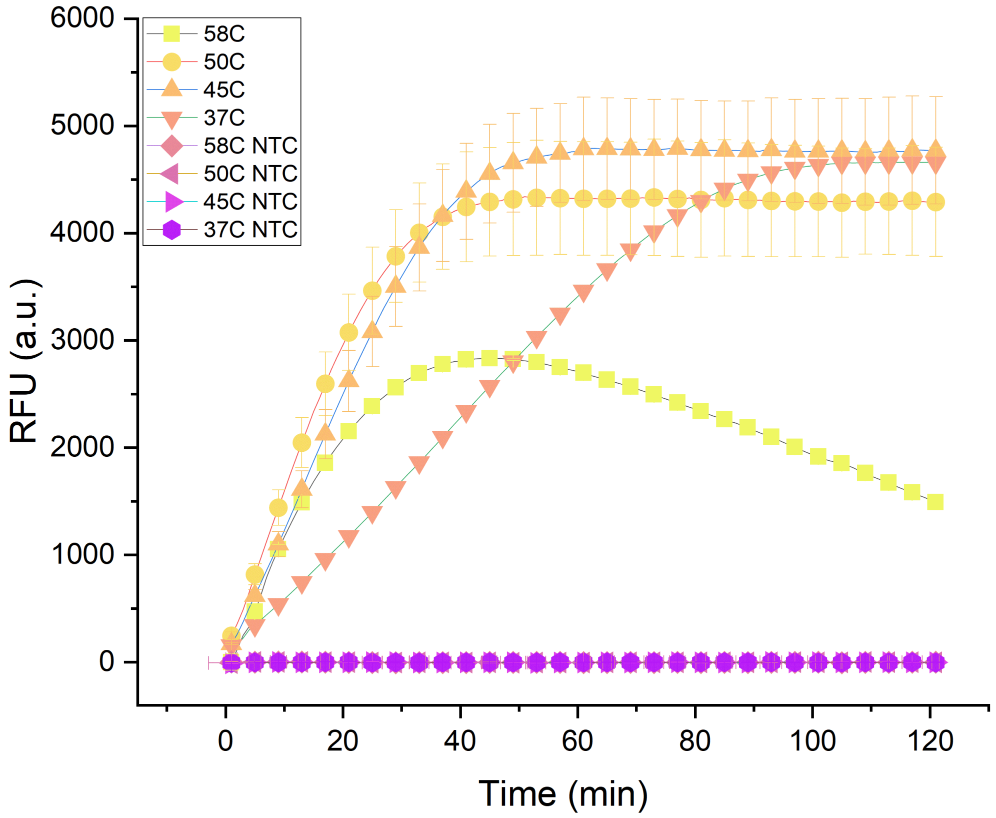


**Figure S4: CRISPR-based detection of synthetic AAV D-sequence.** 50 pM of a short, 42 nt sequence containing the ITR D-sequence as well as minor secondary structure was detected with CRAAVE assay. Enzymatic activity increased from 37C to 45C evidenced by increased slope of temporal fluorescence. Significant additional benefit was not achieved above 45C. All temperatures displayed linear increases in fluorescence beginning at time 0.

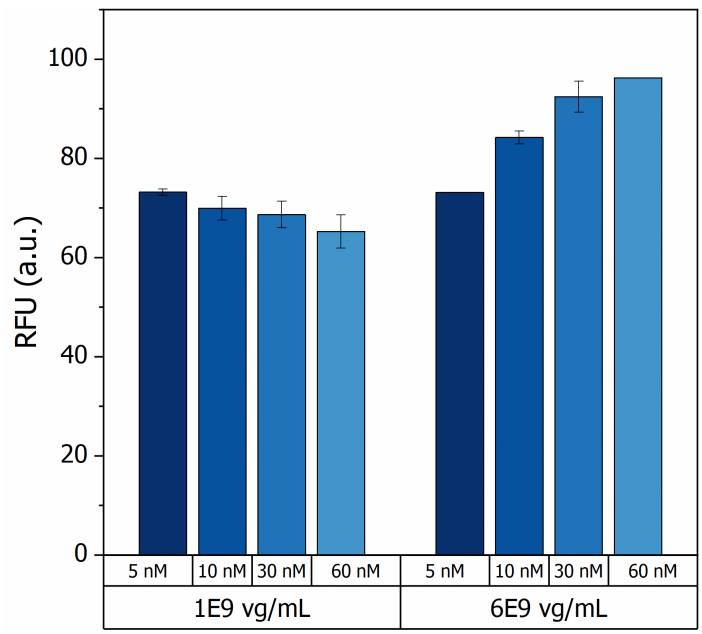


**Figure S5: Assay Signal dependence on Cas RNP concentration**. Virovek rAAV was incubated with varying concentrations of Cas RNP (5 nM – 60 nM) and higher signal was observed at lower rAAV concentrations with lower concentrations of Cas RNP. However, the inverse was observed for relatively higher rAAV concentrations.

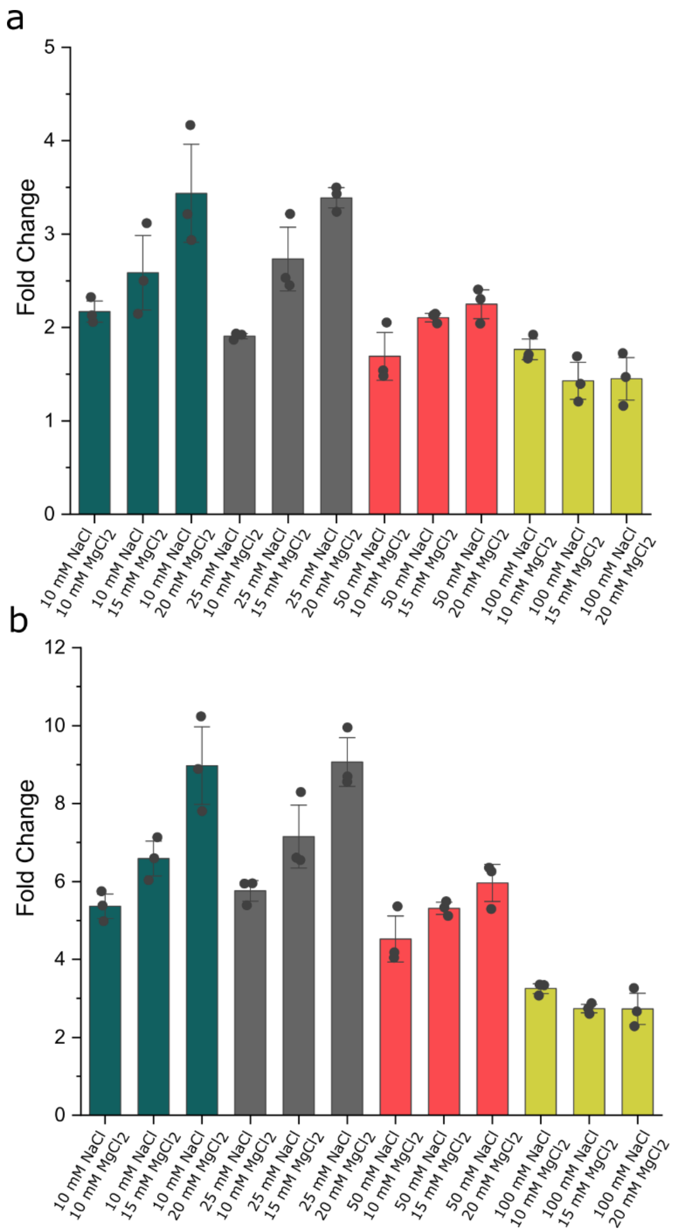


**Figure S6: Ionic Strength Effects on CRAAVE assay fold change.** (a) Fluorescence signal fold changes after 30 minutes of reaction. (b) Fluorescence signal fold changes after 1 hour of reaction.

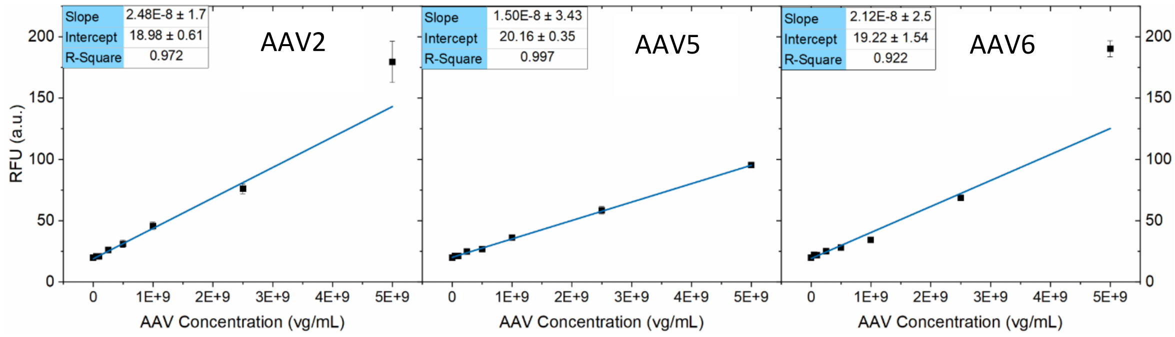


**Figure S7: Standard Curves for CRAAVE assay on various AAVs tested**. Affinity purified AAV2,5, and 6 were serially diluted and assayed with the CRAAVE assay, and standard curves were constructed from fluorescence intensities after 30 minutes. AAV2 encapsidated CellBiolabs pAAV-GFP construct, AAV5 encapsidated Plasmid Factory pAAV-ssGFP, and AAV6 encapsidated a combination of both constructs. The CRAAVE assay was most sensitive to AAV2 and least sensitive to AAV5. The CRAAVE assay displays a median sensitivity toAAV6, likely as it contains a mixture of constructs from AAV2 and 5.

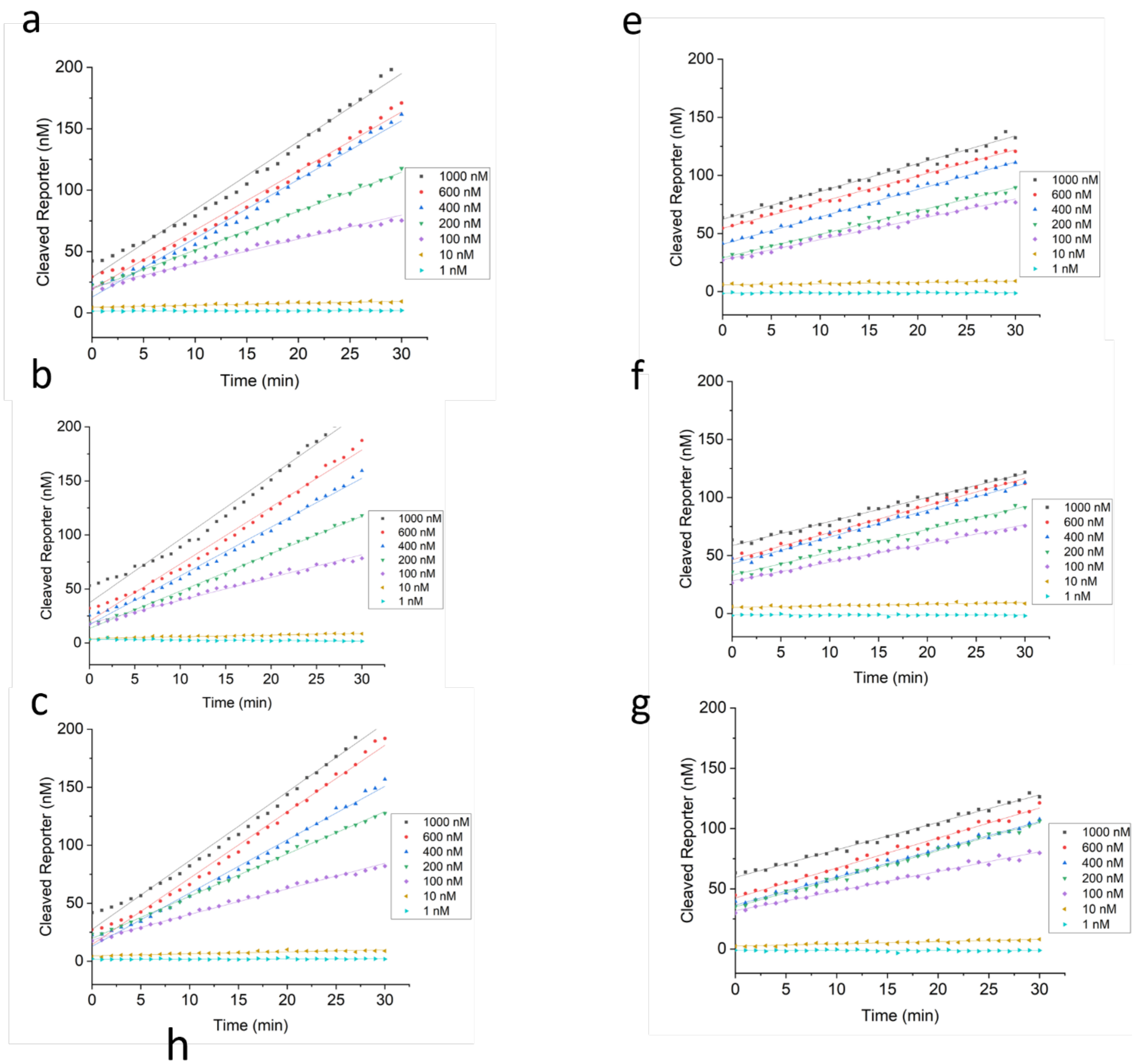


h

| Parameter | Real AAV | Synthetic AAV |
|---|---|---|
| $K_{cat}$ | 19.67±3.08 (1/s) | 6.17±0.60 (1/s) |
| $K_m$ | 485.95 ± 119.04 (nM) | 111.49 ± 41.10 (nM) |
| $V_{max}$ | 9.44 ± 1.48 (nM/min) | 2.96 ± 0.29 (nM/min) |

**Figure S8: Cleaved reporter versus time and Michaelis-Menten kinetic values.** Assay velocities were fitted to Michaelis-Menten kinetics from n=3 replicates of 7 concentrations of cleaved reporter. (a-c) individual kinetic data for cleavage after Cas12a activation with rAAV genomes. (e-g) individual kinetic data for cleavage after Cas12a activation with synthetic AAV DNA. (h) kinetic parameters obtained from fitting kinetic data to the Michaelis-Menten model

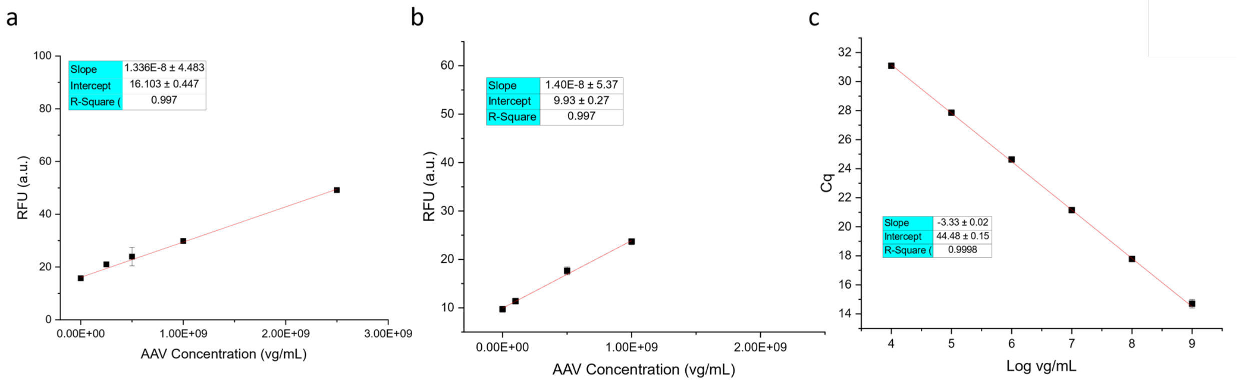


**Figure S9: Assay calibration curves on pAAV-GFP construct.** All calibration curves were constructed with real AAV2 rather than plasmid DNA to prevent assay variation resulting from sample differences between standards and samples. (a) Microplate CRAAVE Assay calibration curve, (b) on-chip CRAAVE calibration curve with removed 5e9 vg/mL data point due to signal saturation, and (c) qPCR calibration curve.

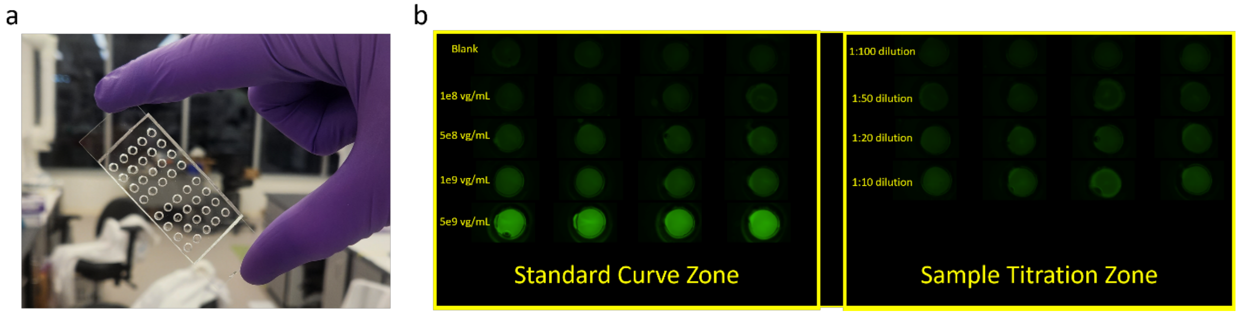


**Figure S10: CRAAVE Assay Chip Design and Readout.** (a) The chip-based CRAAVE assay is constructed with wells punched through PDMS bonded to a glass slide. Lyophilized CRAAVE assay reagents are present in each well requiring only addition of AAV sample and sealing with mineral oil for assay initiation. (b) Stitched image of individual fluorescence micrographs from the CRAAVE assay chip. The left array of wells was used for constructing the standard curve, and the right array of wells was used for sample interrogation.
